# TideCluster: clustering and classification of tandem repeats for comparative and pangenomic analysis

**DOI:** 10.64898/2026.11.06.764801

**Authors:** Petr Novák, Jiří Macas

## Abstract

Satellite DNA, composed of tandem arrays of relatively short repeated units, is a ubiquitous component of plant and animal genomes. Despite its biological importance, systematic investigation of satellite DNA remains challenging, in part because integrated bioinformatic tools for structured comparative analyses across individual arrays, chromosomes, and genomes are lacking. Here, we present TideCluster, a reference-independent pipeline that converts fragmented tandem repeat detections into genome-wide, interpretable repeat family catalogs. Starting from a genome assembly, TideCluster consolidates detected regions into arrays, clusters arrays by sequence similarity, reconstructs representative monomer consensus sequences, estimates locus-specific periodicity, identifies higher-order repeats, and generates GFF3 annotations. In addition to predefined visualizations, including the chromosomal distributions of satellite repeat families, TideCluster produces information-rich outputs that can be adapted for user-specific downstream analyses. We demonstrate its capabilities using the well-characterized genome of *Arabidopsis thaliana* and a set of Solanum species with highly diverse satellite repeat complements that vary markedly in abundance and have monomer lengths ranging from a few base pairs to more than 20 kb.

**Availability & Implementation:** TideCluster is freely available at https://github.com/kavonrtep/TideCluster under the GNU General Public License v3; test data and analysis scripts are deposited at Zenodo (doi:10.5281/zenodo.22659516).

## Statement of Need

Tandem repeats represent a structurally distinct class of repetitive DNA characterized by the head- to-tail arrangement of their repeated units, also called monomers. Although tandem repeats in eukaryotic genomes are highly diverse in structure, those forming long, uninterrupted arrays with monomers ranging from tens to thousands of nucleotides—collectively termed satellite DNA (satDNA)—are predominant [1]. SatDNA constitutes a substantial and highly dynamic fraction of eukaryotic genomes, forming functional chromosomal domains such as centromeres and telomeres and contributing to genome size and structural variation [1–4].

Despite their biological importance, the systematic investigation of satellite DNA in complex eukaryotic genomes has lagged behind the characterization of other genomic components. This delay has largely been driven by the inherent difficulty of assembling highly repetitive genomic regions enriched in satDNA [3,5], as well as by the historical lack of appropriate bioinformatic tools capable of resolving such sequences. Consequently, first-generation approaches, such as TAREAN [6] and related methods, were primarily designed to analyze unassembled short-read shotgun data. However, analyses based on unassembled reads cannot capture the full sequence diversity of satellite repeats nor provide locus-specific information about individual tandem repeat arrays, which is essential for understanding the mechanisms of satDNA evolution. Such information becomes accessible only in the context of genome assemblies. Fortunately, recent advances in long-read sequencing technologies have led to a rapid increase in the availability of high-quality genomic resources, including telomere-to-telomere (T2T) assemblies [4,5], thereby enabling more comprehensive analyses of repetitive regions and underscoring the need for dedicated bioinformatic tools capable of systematically characterizing satDNA at the level of assembled genomes.

Characterization of satellite DNA in genome assemblies should ideally satisfy several key requirements: (1) detection of tandem repeat arrays based solely on sequence structure, without relying on predefined reference libraries; (2) inference of monomer length for each array and identification of potential higher-order repeat (HOR) organization; (3) genome-wide classification of detected arrays into satDNA families based on sequence similarity; and (4) support for comparative analysis of satDNA composition across species. In addition, such analyses must be computationally efficient to enable application to large and repeat-rich genomes. To the best of our knowledge, no existing tool fulfills all of these criteria simultaneously. Many approaches, particularly those developed for centromeric satellite analysis in model organisms (e.g., HORmon [7] and HiCAT [8]), rely on reference sequences for detection and characterization of satellite arrays or are restricted to strictly defined short centromeric monomers of approximately 100–200 bp (e.g., centroAnno [9] and CENdetectHOR [10]). More general structural detection of tandem repeat arrays is provided by tools such as Tandem Repeats Finder (TRF) [11], ULTRA [12], TRASH [13], and TideHunter [14], but these methods typically treat each detected array as an independent unit without higher-level integration. Consequently, no existing framework provides a unified solution for hierarchical characterization of tandem repeats within genomes and systematic comparison across multiple species.

To address this gap, we present TideCluster, a bioinformatics pipeline that converts fragmented array-level detections of tandem repeats into interpretable repeat-family catalogs that can be compared within and across assemblies. We demonstrate TideCluster’s capabilities using well-characterized genomes, in which recovery of known families can be assessed, including the model species *Arabidopsis thaliana [4]* and a set of Solanum species known for diverse satellite DNA landscapes that differ in sequence composition, monomer length, and abundance [15–18].

## Implementation

TideCluster is a computational pipeline implemented in Python [19] and R [20], executed from the command line, with a genome assembly in FASTA format as its only required input. It uses TideHunter [14] to identify genomic regions composed of tandemly repeated sequences, which are then processed through the series of steps outlined in Figure 1. First, TideHunter-reported regions are consolidated into tandem repeat arrays (TRAs), which are subsequently subjected to similarity-based clustering to form tandem repeat clusters (TRCs) corresponding to repeat families. For each sufficiently abundant TRC, TideCluster reconstructs a representative consensus sequence, annotates all constituent TRAs with the corresponding TRC label, and characterizes intra-array variation, including monomer-length diversity and the presence of higher-order repeats (HORs).

**Figure 1.**
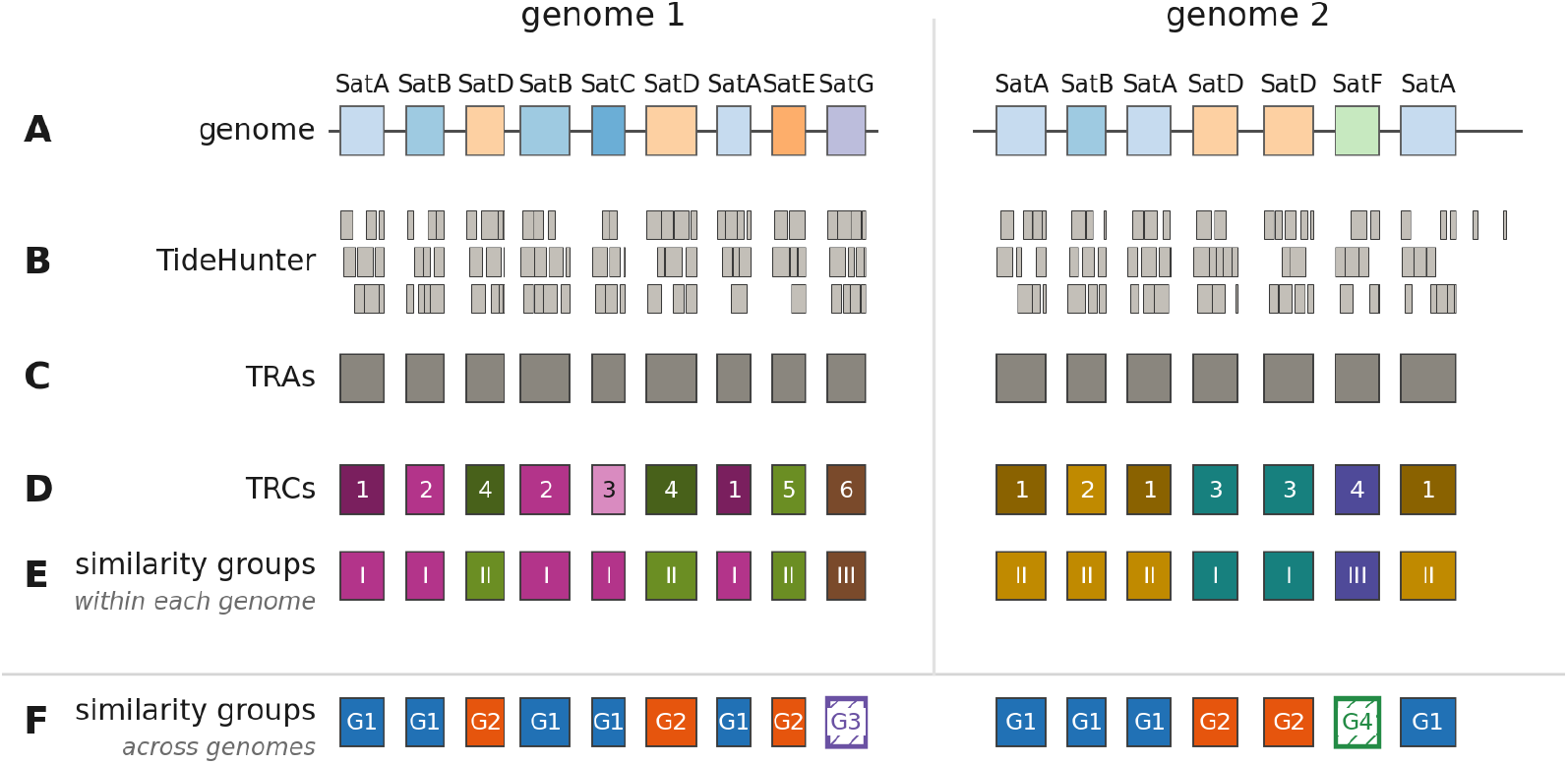
TideCluster workflow illustrated for two hypothetical genomes. (A) Tandem-repeat arrays in the input genomes. (B) Fragmented and overlapping TideHunter detections. (C) Merged tandem-repeat arrays (TRAs). (D) Similarity-based tandem-repeat clusters (TRCs). (E) Within-genome groups of related TRCs. (F) Cross-genome TRC groups. Colors denote sequence relationships; shading distinguishes families within a related group. Hatched groups in F occur in only one genome. Outputs are summarized in Table 1.

In addition, TRCs can optionally be compared against user-provided repeat databases or against TRCs identified in other species to define cross-genome TRC groups. These groupings are based on explicit, user-adjustable sequence similarity criteria and should therefore be interpreted as operational classifications rather than fixed biological categories. The pipeline can be run as a complete workflow or in individual steps. The final output consists of GFF3-formatted annotations of satellite DNA arrays in the assembly, together with additional data intended to support downstream analyses (Table 1).

**Table 1.** Principal outputs of the TideCluster pipeline. Letters correspond to the stages in Figure 1.

|  | Step | Output | Contents |
|---|---|---|---|
| **A** | Input | — | Genome assembly |
| **B** | Detection | tc_tidehunter.gff3 | Tandem repeat regions |
| **C + D** | Merging and clustering | tc_clustering.gff3 | Consolidated tandem arrays assigned to tandem repeat clusters (TRCs) |
| **D** | Consensus | Consensus FASTA (monomer and dimer) | Cluster (TRC) monomer consensus |
|  | Annotation *(optional)* | tc_annotation.gff3,<br>tc_annotation.tsv | TRCs annotated with custom database |
| **E** | Within-genome grouping | tc_trc_superfamilies.csv | The TRC similarity groups assignment |
| **F** | Comparative analysis *(optional)* | Cross-genome group table; updated GFF3 per genome | Cross-genome similarity groups assignment |
| **A–F** | Report | tc_index.html | Comprehensive HTML report (Monomer length,higher-order repeats, SSR and rDNA) |

### Tandem repeat detection

Tandem repeat arrays (TRAs) are identified by parsing the output of TideHunter executed with parameters targeting repeats with monomer lengths between 40 and 3000 bp. Optionally, the analysis can be extended to include tandem repeats with monomers up to 25,000 bp long, enabling detection of features such as rDNA arrays. The minimum number of tandem repetitions and the maximum sequence divergence of monomers in reported arrays are set to 5 and 0.25, respectively, but these values, as well as most other parameters, are user-adjustable. Because TideHunter’s performance degrades with increasing sequence length, chromosome-scale or long-contig assemblies are partitioned into overlapping 50 Mbp chunks for this step, and coordinates are subsequently restored to the original genomic frame. By design, TideHunter reports a single tandem repeat array as several adjacent, often overlapping regions rather than as a single feature (Figure 1B); TideCluster therefore converts this fragmented output into coherent arrays (Figure 1C). Subsequent analysis is performed only on TRAs that exceed a defined minimum length (5000 bp by default).

### Sequence similarity-based clustering of tandem repeat arrays

This step aims to detect significant sequence similarities among individual TRAs identified in the assembly and assign them to tandem repeat clusters (TRCs) that broadly correspond to satDNA families (Figure 1D). It uses a two-stage procedure designed to balance computational efficiency and sensitivity. First, the TideHunter consensus sequences representing each TRA are grouped with MMseqs2 [21] to reduce redundancy among nearly identical repeats. Representatives of the MMseqs2 groups are then subjected to a more sensitive all-against-all BLASTN [22] comparison to detect similarities of at least 75% spanning at least 80% of the shorter of the two sequences. Consensus monomer sequences exceeding this similarity threshold are linked by single linkage clustering [23] to form a single TRC. TRCs are then ranked by total genomic coverage and numbered TRC_1, TRC_2, … from largest to smallest.

To infer genome-wide monomer consensus sequences and their most frequent variants for each abundant satDNA family, TRCs whose combined TRA length exceeds a minimum threshold (50 kb by default) are subjected to TAREAN analysis [6]. All TRA regions assigned to a given TRC are decomposed into k-mers, and overall k-mer frequencies within the TRC are calculated. TAREAN then constructs a de Bruijn graph from the most frequent overlapping k-mers and identifies circular paths in the graph corresponding to candidate monomer variants. The path with the strongest k-mer support is selected as the consensus monomer, while other top-ranked paths are recovered as its major variants. For a detailed description of the reconstruction algorithm, see [6].

Because the described procedure for clustering and consensus reconstruction cannot be efficiently applied to low-complexity tandem repeats, such as telomeric repeats and simple sequence repeats (SSRs), these regions are analyzed differently: for each TRA that consists of more than 90% low-complexity sequence, as assessed by the NCBI dustmasker utility [24], the underlying short motif is reconstructed and converted to a canonical form by selecting the lexicographically smallest sequence among all cyclic rotations of the motif and its reverse complement. This normalization converts motifs that differ only in their starting position or strand orientation to the same SSR motif. TRAs sharing the same SSR motif are then placed into the same TRC group.

### Periodicity analysis and HOR detection

In addition to inferring a consensus monomer sequence and its major variants globally for each abundant TRC, TideCluster also analyzes the periodicity of individual TRAs to reveal locus-specific variation in monomer length. It also detects higher-order repeats (HORs), defined as tandemly repeated units composed of a characteristic, ordered combination of two or more monomer variants. This procedure exploits the regular spacing of shared k-mer patterns along tandem arrays to infer underlying repeat structure in a reference-free manner (Figure 2). TideCluster derives candidate periodicities corresponding to potential tiling units (including both monomers and their higher-order multiples) from k-mer recurrence distances along each array (Figure 2A-B). Each candidate is scored by the median sequence identity between adjacent repeats, and the highest-scoring one is selected as the strongest period, i.e., the unit that best tiles the array (Figure 2C). The monomer size is then defined as the shortest unit that divides this strongest period while still maintaining sufficient tiling identity. In simple arrays, the strongest period corresponds to the monomer; in higher-order arrays, it corresponds to a composite unit made of multiple monomers. The number of monomers per strongest period defines the multiplicity (≥2 for HORs), and TideCluster reports a HOR identity gain as the difference in adjacent-unit identity between the strongest period and the monomer, quantifying the strength of higher-order organization. This analysis requires no prior template or annotation and serves as an exploratory measure of periodic structure alongside monomer estimation.

**Figure 2.**
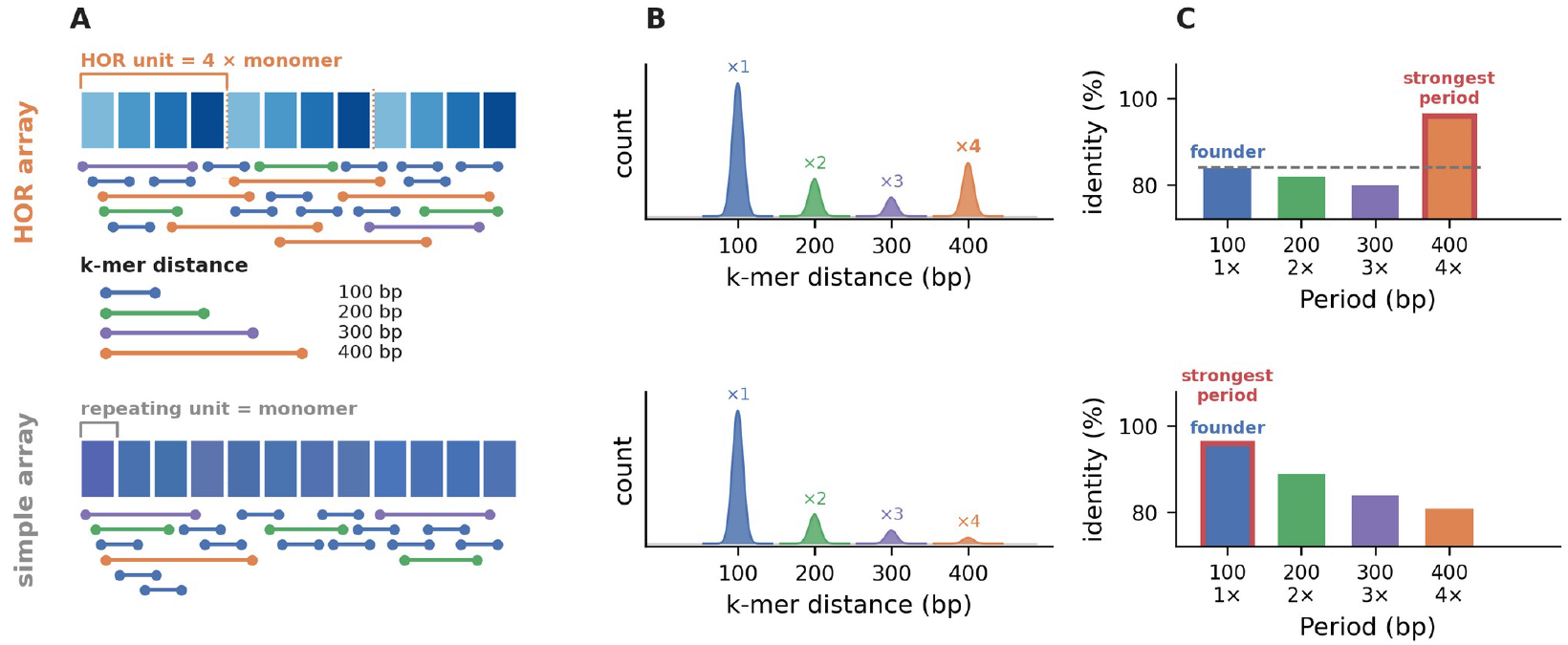
Identification of higher-order repeats in arrays with (top) and without (bottom) higher-order organization. (A) Recurrence distances of identical k-mers along each array. (B) Distance periodograms identifying candidate repeat periods. (C) Mean sequence similarity between adjacent units for each candidate period. The highest-scoring period (red outline) comprises four monomers in the higher-order array and one monomer in the simple array. HOR identity gain is the improvement over the monomer-period identity.

### Similarity-based annotation and grouping of TRCs

After TRCs and their representative consensus sequences have been established, TideCluster provides three complementary similarity-based analyses for organizing and interpreting the identified repeat families. These consist of (1) grouping sequence-related TRCs within an individual genome into TRC groups, (2) annotating TRCs by comparison with known repeats, and (3) identifying similar TRCs across multiple genomes or samples as cross-genome TRC groups. These analyses add relationships and annotations to the results and should be interpreted as operational classifications rather than formal evolutionary or taxonomic categories.

To reveal broader relationships among repeat families within a genome, TideCluster compares the TAREAN-derived consensus sequences of TRCs pairwise using BLAST. TRCs whose consensus sequences exceed a configurable similarity-score threshold are grouped into TRC groups (Figure 1E). Because this criterion permits partial sequence similarity, a TRC group may include TRCs with different monomer lengths or TRCs that share only a part of their consensus sequences.
TideCluster can optionally annotate TRCs against a user-supplied library of known tandem repeats. Consensus sequences associated with each TRC are analyzed with RepeatMasker [25], and similarity matches are added to the GFF3 output and summarized at the TRC level. TRCs without a supported library match remain unannotated.
For comparative analyses, consensus sequences from TRCs identified independently in multiple genomes or samples are combined and compared using MMseqs2. Sequence pairs satisfying configurable identity and alignment-coverage thresholds form a similarity graph, which is partitioned by a community-detection algorithm [26] into cross-genome TRC groups (Figure 1F). The analysis preserves the original sample-specific TRC identifiers and integrates their annotations and genomic abundance, thereby enabling the distribution and relative abundance of related tandem repeat families to be compared across samples.

## Results

To evaluate TideCluster across genomes differing in size, repeat content, and assembly quality, we applied it to the reference genome of *Arabidopsis thaliana* and eleven publicly available *Solanum* assemblies from the tomato and potato clades (Table 2). The well-characterized centromeric and pericentromeric satellites of *A. thaliana* [4,27] provided a reference for evaluating repeat recovery, clustering, and annotation, whereas the satellite-rich *Solanum* genomes tested the pipeline on larger genomes containing diverse repeats, including families with long monomers [15,16,18,28,29]. Together, these datasets were used to assess the biological validity of the inferred repeat hierarchy, computational scalability, and comparative analysis across genomes.

**Table 2.** Investigated species and overview of TideCluster processing of tandem-repeat regions detected by TideHunter, including the number of instances at each stage from initial detection through array construction, clustering, and TRC similarity grouping. The table also reports the total amount of detected tandem repeats (TR), the range of monomer lengths, and the proportion of TRAs containing higher-order repeats. The TideCluster workflow is illustrated in Figure 1.

| Species | Accession | Assembly (Mb) | TideHunter regions | TRAs | TRCs | TRC similarity groups | TR (Mb) | TR (% of genome) | Monomer range (bp) | TRAs with HOR (%) |
|---|---|---|---|---|---|---|---|---|---|---|
| *Arabidopsis thaliana* | GCA_028009825.2 | 142.48 | 2,427 | 154 | 28 | 17 | 22.14 | 15.54 | 97–13564 | 85 (55%) |
| *Solanum chilense* | GCA_032847665.1 | 901.27 | 9,284 | 105 | 59 | 59 | 1.18 | 0.13 | 3–7111 | 58 (55%) |
| *Solanum galapagense* | GCA_025758845.1 | 806.40 | 10,348 | 489 | 76 | 71 | 8.55 | 1.06 | 3–9779 | 189 (39%) |
| *Solanum lycopersicum* | GCA_036512215.2 | 832.77 | 20,820 | 747 | 73 | 66 | 40.75 | 4.89 | 3–16326 | 360 (48%) |
| *Solanum lycopersicum var. cerasiforme* | GCA_025759795.1 | 803.62 | 14,789 | 648 | 77 | 61 | 13.39 | 1.67 | 3–18088 | 316 (49%) |
| *Solanum pennellii* | GCA_001406875.2 | 926.58 | 9,627 | 354 | 87 | 86 | 4.00 | 0.43 | 2–4872 | 171 (48%) |
| *Solanum pimpinellifolium* | GCA_034621305.1 | 833.19 | 12,598 | 434 | 66 | 63 | 7.57 | 0.91 | 1–11954 | 194 (45%) |
| *Solanum acaule* | GCA_046563305.1 | 695.56 | 10,124 | 589 | 141 | 126 | 16.60 | 2.39 | 3–23666 | 282 (48%) |
| *Solanum commersonii* | GCA_018258275.1 | 731.62 | 8,253 | 190 | 70 | 70 | 2.09 | 0.29 | 7–9000 | 88 (46%) |
| *Solanum microdontum* | GCA_054051385.1 | 771.71 | 10,491 | 459 | 123 | 107 | 25.06 | 3.25 | 6–9250 | 234 (51%) |
| *Solanum tuberosum* | GCA_015076265.1 | 716.17 | 9,728 | 511 | 159 | 135 | 13.24 | 1.85 | 3–11157 | 208 (41%) |
| *Solanum verrucosum* | GCA_031230405.1 | 683.98 | 9,497 | 500 | 133 | 115 | 13.52 | 1.98 | 7–14011 | 236 (47%) |

In all tested assemblies, thousands of tandemly organized sequence regions were identified by TideHunter and progressively consolidated by TideCluster into decreasing numbers of contiguous arrays, TRCs, and TRC similarity groups, as summarized in Table 2. Tandem-repeat content ranged from 0.13% to 4.89% among the *Solanum* assemblies and reached 15.54% in *A. thaliana*, owing to its small genome size and extensive CEN180 and rDNA arrays. Consensus monomer lengths ranged from a few base pairs to more than 23 kb. The monomers were found to form HORs in a substantial proportion of tandem-repeat arrays (Table 2). All assemblies were analyzed on a single workstation using moderate memory, demonstrating the practical scalability of the approach to gigabase-sized plant genomes (Table 3).

**Table 3.** Computational performance of single-genome analyses using 16 threads on an AMD Ryzen 7 5700X workstation with 128 GB RAM.

| Species | Assembly (Mb) | Wall time (hh:mm:ss) | Peak RAM (GB) |
|---|---|---|---|
| *Arabidopsis thaliana* | 142.48 | 00:18:54 | 22.1 |
| *Solanum verrucosum* | 683.98 | 01:05:19 | 11.3 |
| *Solanum acaule* | 695.56 | 00:58:52 | 11.6 |
| *Solanum tuberosum* | 716.17 | 02:03:35 | 12.5 |
| *Solanum commersonii* | 731.62 | 00:43:11 | 8.4 |
| *Solanum microdontum* | 771.71 | 00:56:42 | 11.6 |
| *Solanum lycopersicum var. cerasiforme* | 803.62 | 01:29:15 | 10.8 |
| *Solanum galapagense* | 806.40 | 00:45:13 | 8.5 |
| *Solanum lycopersicum* | 832.77 | 01:35:25 | 27.9 |
| *Solanum pimpinellifolium* | 833.19 | 00:51:00 | 10.1 |
| *Solanum chilense* | 901.27 | 00:45:25 | 11.1 |
| *Solanum pennellii* | 926.58 | 00:51:02 | 9.5 |

In *A. thaliana*, TideCluster recovered the five expected major tandem-repeat families, including the satellite repeats CEN180, AthSat500, and AthSat160, as well as the 45S and 5S ribosomal DNAs (Figure 3). These five families accounted for 21.5 Mb, or 97.2% of the recovered tandem-repeat sequence, and were distributed among 14 of the 28 TRCs (Table 4). The remaining 14 TRCs contained only 2.8% of the recovered sequence and represented small, low-abundance arrays without a well-supported annotation. TRC similarity grouping merged CEN180 and AthSat500 into one group (G1, Table 4), while the remaining TRC similarity groups included a single repeat family. This is consistent with the known sequence similarity between part of the AthSat500 monomer and CEN180 [27]. The HOR analysis revealed that most CEN180 arrays (84.7%) have higher-order organization, consistent with [4], who reported 95.5% of CEN180 as being composed of higher-order repeats. The discrepancy between the two estimates is most likely due to stricter criteria for HOR identification in TideCluster than those used in [4].

**Table 4.** Recovery of known *Arabidopsis thaliana* tandem-repeat families. Families were assigned when a reference library sequence covered at least 50% of the TRC consensus; all other TRCs were classified as unannotated. TR length is the total genomic length of the corresponding arrays. TRAs, tandem-repeat arrays; TRCs, tandem-repeat clusters.

| Annotated family | Class | Library accession | TideHunter regions | TRAs | TRCs | Similarity group ID | TR length (bp) | Genome proportion (%) |
|---|---|---|---|---|---|---|---|---|
| CEN180 | satellite | X04322.1 | 611 | 72 | 1 | G1 | 11,421,836 | 8.02 |
| AthSat500 | satellite | X06471.1 | 18 | 10 | 3 | G1 | 230,828 | 0.16 |
| AthSat160 | satellite | X06472.1 | 47 | 7 | 1 | G2 | 201,194 | 0.14 |
| 45S rDNA | rDNA | X52322.1 | 947 | 35 | 7 | G3 | 7,722,411 | 5.42 |
| 5S rDNA | rDNA | M65137.1 | 50 | 13 | 2 | G4 | 1,935,817 | 1.36 |
| Unannotated | <50% / none | — | 43 | 17 | 14 | G5-G17 | 629,786 | 0.44 |

**Figure 3.**
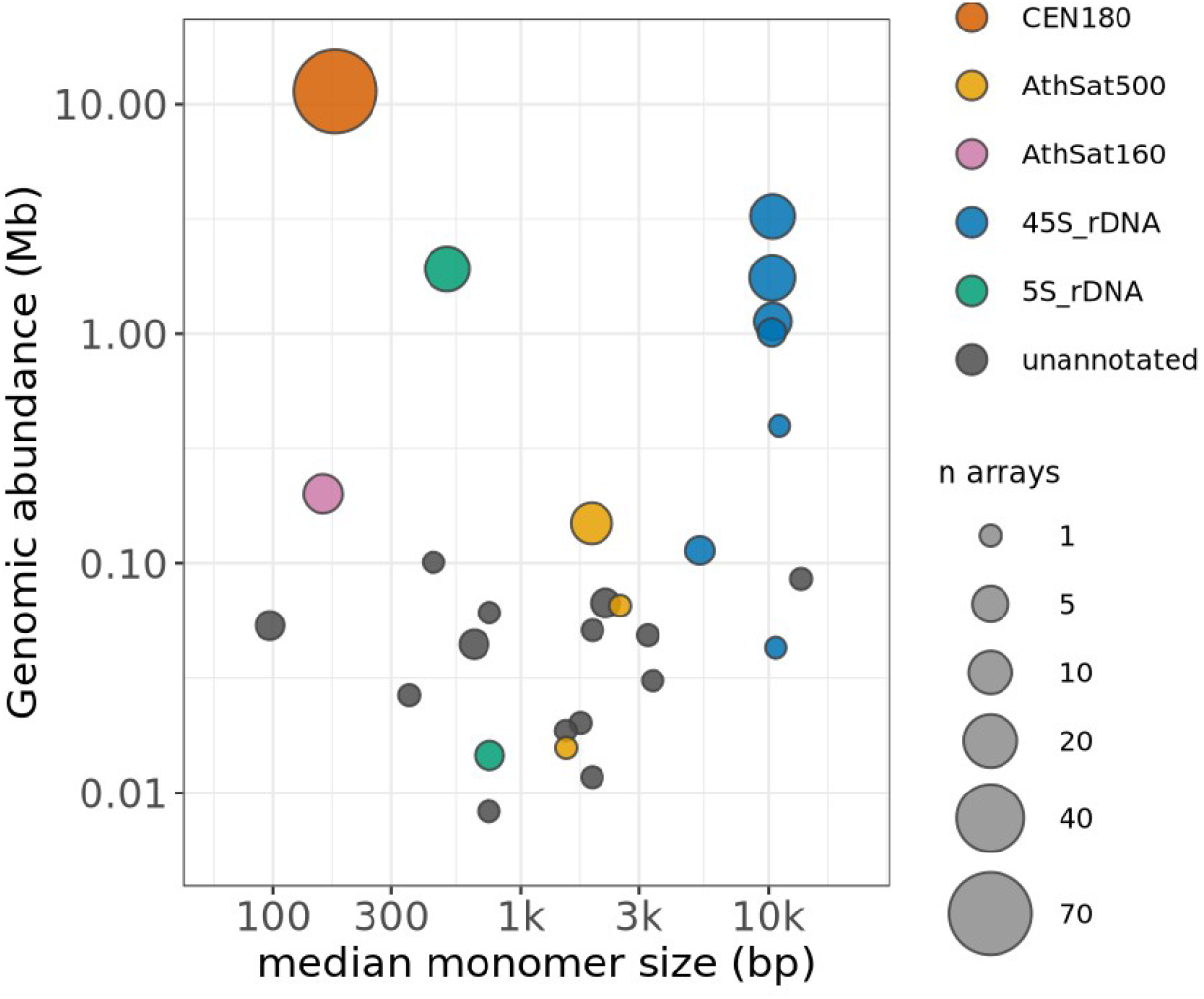
Tandem-repeat cluster (TRC) landscape of *Arabidopsis thaliana*. Each point represents one TRC. The axes show median founder-monomer length and total genomic abundance; point size indicates the number of arrays (TRAs). Both axes are logarithmic. Colors identify repeat families assigned by significant similarity to sequences in the user-provided reference library of known repeats, defined as coverage of at least 50% of the TRC consensus; unannotated TRCs are gray.

Application of TideCluster to the selected *Solanum* assemblies recovered the major satellite DNA families previously reported from these species (Figure 4), despite substantial variation in monomer length, including repeats with monomers up to 23 kb that are not detectable by most alternative tools. In addition to characterizing the tandem-repeat composition of individual genomes, TideCluster provided chromosome-level distributions of individual repeat families (Figure 5). These distributions were consistent with published sequence and cytogenetic data: TGRI/pSB7 [30,31] and SPG/CL34 [17,32] were among the most abundant TRCs across the analyzed genomes, while the strictly terminal localization of SPG/CL34 and centromeric location of St49 [16] and Sv44 [18] agreed with their reported cytogenetic distribution.

**Figure 4.**
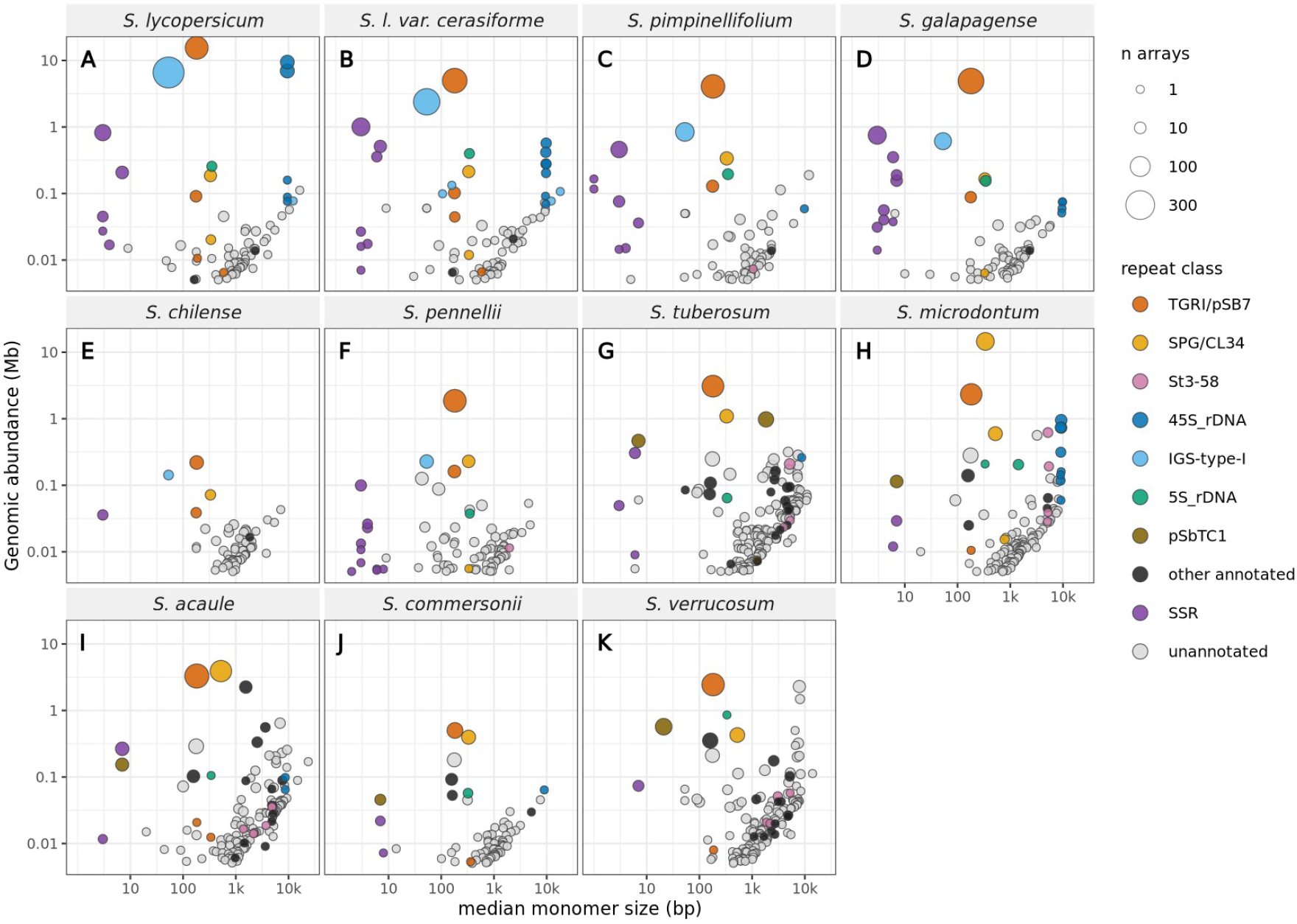
Tandem-repeat cluster landscapes of eleven *Solanum* genomes: tomato clade (**A–F**) and potato clade (**G–K**). Axes and point sizes are defined as in Figure 3. Colors identify selected published repeat families, rDNA, simple-sequence repeats (SSRs), and other library-annotated TRCs; unannotated TRCs are gray.

**Figure 5.**
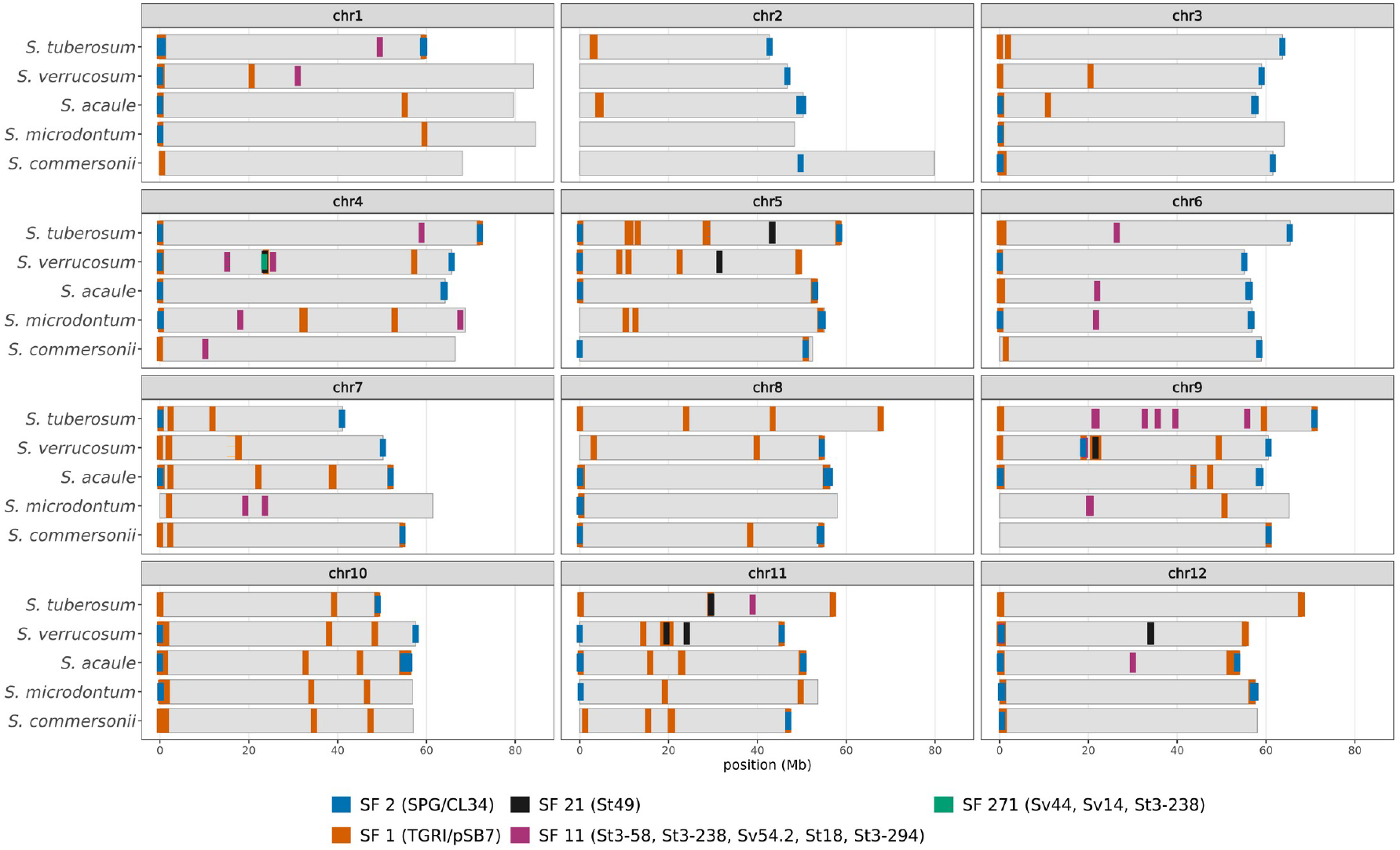
Chromosomal distributions of previously described satellite repeats in five potato-clade Solanum genomes. Panels group homeologous chromosomes 1–12 from the examined species. Gray bars represent chromosomes, and tick colors denote different cross-genome similarity groups (SF1, SF2, SF11, SF21, and SF271), each comprising previously published repeats as specified in the graphical legend.

Although known repeats shared among species can be readily identified using a reference library (Figure 5), the TideCluster comparative-analysis module provides a more comprehensive, reference-independent assessment of shared and species-specific tandem repeats. As shown in Figure 6, it identifies interspecific similarity groups comprising TRCs with significant sequence similarity in at least two analyzed species, including newly characterized repeats that are absent from existing libraries. Reference-library annotations are nevertheless retained to facilitate the identification of known satellite families. Consistent with Figure 5, the TGRI/pSB7 and SPG/CL34 repeats were detected across all analyzed genomes, together with previously uncharacterized groups such as SF10. The analysis also separated the genomes into the tomato and potato phylogenetic clades, each characterized by a distinct subset of shared tandem repeats.

**Figure 6.**
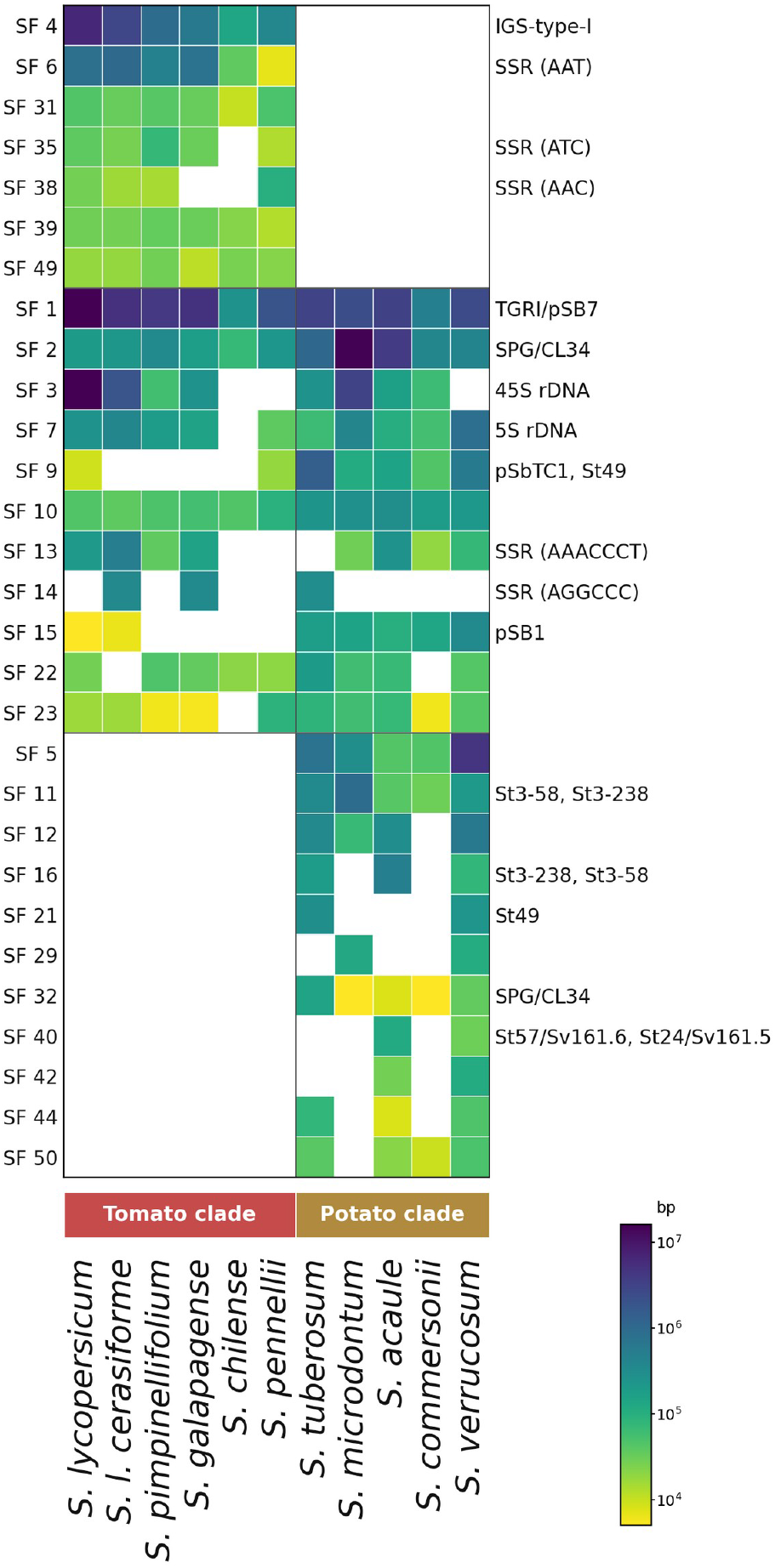
Distribution and abundance of the 29 largest cross-genome TRC similarity groups across eleven *Solanum* genomes. Columns are genomes grouped into tomato and potato clades; rows are TRC groups classified as tomato-clade-specific, shared between clades, or potato-clade-specific. Cell color represents log10 of the total length of the corresponding repeat sequence in each genome; blank cells indicate no detected group member. Similarities to repeats from reference library are shown at right.

## Availability of Source Code and Requirements

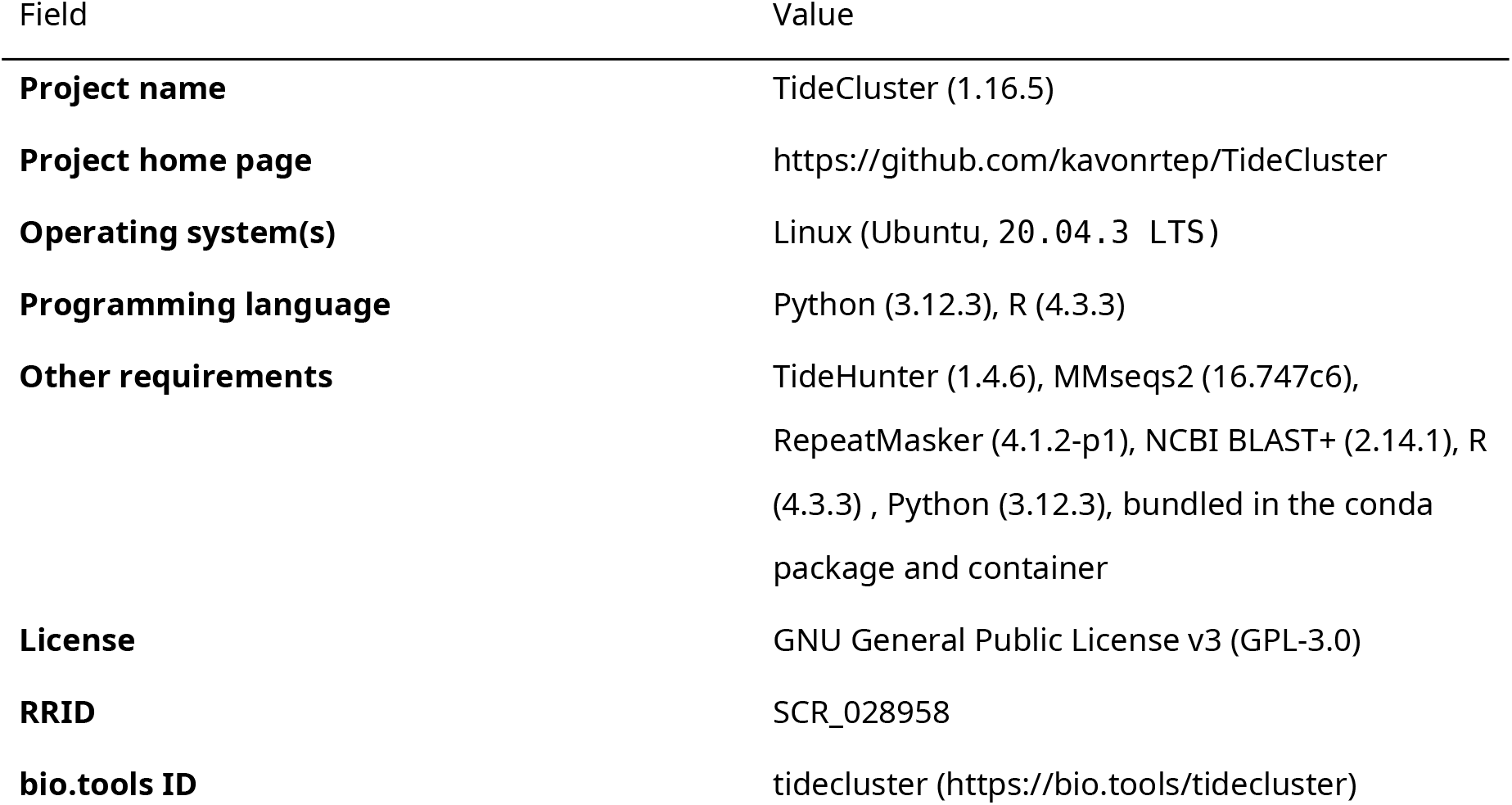

### Data availability

The supporting data are available in Zenodo [33]. They include full TideCluster output for all species analyzed and the library of tandem repeats used for annotation.

## Abbreviations

HOR: higher-order repeat
SSR: simple sequence repeat
TR: tandem repeat
TRA: tandem repeat array
TRC: tandem repeat cluster.

## Declarations

### Ethical Approval

Not applicable.

### Authors’ Contributions

Contributions are described using the NISO CRediT taxonomy. Petr Novák: Conceptualization, Data curation, Formal analysis, Investigation, Methodology, Software, Validation, Visualization, Writing – original draft, Writing – review & editing. Jirí Macas: Conceptualization, Funding acquisition, Investigation, Methodology, Project administration, Resources, Supervision, Validation, Writing – original draft, Writing – review & editing.

### Disclosure of Use of AI-Assisted Tools Including Generative AI

The authors used the generative AI assistant Claude (Anthropic), accessed through the Claude Code interface, for data-analysis and figure scripting and for drafting and editing the manuscript [34]. Claude Opus 4.8 (30 May to 8 September 2026) and Claude Opus 5 (11 to 14 August 2026) were used by P.N. No figures were generated or altered by generative AI. All AI-assisted output was verified and edited by the authors, who take full responsibility for the content; AI tools do not meet authorship criteria and are not listed as authors.

## Acknowledgments

The authors acknowledge the ELIXIR CZ research infrastructure (MŠMT Grant No. LM2023055) for support.

